# Environment-conditioned male fertility of HD-ZIP IV transcription factor mutant *ocl4*: impact on 21-nt phasiRNA accumulation in pre-meiotic maize anthers

**DOI:** 10.1101/2020.06.30.180398

**Authors:** Pranjal Yadava, Saleh Tamim, Han Zhang, Chong Teng, Xue Zhou, Blake C. Meyers, Virginia Walbot

## Abstract

Environment-conditioned genic male sterility is a key strategy used to produce hybrid seeds efficiently in many crops, with the exception of maize. The underlying molecular mechanisms of environment-conditioned sterility are poorly understood. Here, we report a derivative line of the male sterile *outer cell layer 4* (*ocl4*) mutant of maize, in which fertility was restored and perpetuated over several generations, under warm growing conditions. Conditionally fertile *ocl4* anthers exhibit the anatomical abnormality of a partially duplicated endothecial layer, just like their sterile counterparts. We profiled the dynamics of phased, small interfering RNAs (phasiRNAs) during pre-meiotic development in fully sterile and various grades of semi-fertile *ocl4* anthers. We found that the biogenesis of 21-nt phasiRNAs is largely dependent on *Ocl4* at three key steps: (1) production of *PHAS* precursor transcripts, (2) expression of miR2118 that modulates precursor processing, and (3) accumulation of 21-nt phasiRNAs. We propose that 21-nt reproductive phasiRNAs buffer development under unfavorable environmental conditions and are dispensable under favorable conditions.

## Introduction

The maize genome encodes 17 homeodomain leucine zipper type IV (HD-ZIP IV) proteins, transcripts of which preferentially accumulate in the epidermis (Sosso et al. 2010; Javelle et al. 2011). Nine family members are expressed in immature tassels: four types are encoded by duplicated genes while *Outer cell layer 4* (*Ocl4*) is a single-copy gene (Javelle et al. 2011). *Ocl4* is expressed in diverse epidermal cell types in multiple organs in addition to the tassel (Javelle et al. 2011). The *ocl4-1* mutant allele is disrupted by a *Mu8* insertion in exon 6 and the plants are male-sterile; mutant plants accumulate fewer transcripts than normal, some transcripts are alternatively spliced, and *ocl4-1* is predicted to encode a truncated partial or complete loss-of-function protein (Vernoud et al. 2009).

During fertile male development, pre-meiotic archesporial cells are specified centrally in each of the four anther lobes. These cells secrete MAC1 protein, a signal that triggers somatic differentiation of neighboring cells forming a ring of primary parietal cells (PPC); these somatic cells then divide periclinally to produce the subepidermal endothecium and the secondary parietal cells adjacent to the archesporial cells (Wang et al. 2012). This division also marks an exit from pluripotency in the soma (Kelliher and Walbot 2014). In *ocl4* mutants, the earliest steps of anther development are normal; however, lobe architecture deviates after the PPC periclinal division. In the outer hemisphere of each lobe, underlying the epidermal domain of *Ocl4* expression, endothecial cells undergo an ectopic periclinal division producing a bilayer (Vernoud et al. 2009). The bilayer defect is cytologically complete about the third day of maize anther development (0.4 mm anther length), and it has been proposed that epidermal cells produce a signal to repress endothecial periclinal cell division (Wang et al. 2012). Generally, *ocl4-1* plants are male-sterile but were reported to show sporadic partial fertility under greenhouse growing conditions (24 °C /19 °C and 16 h light/ 8 h dark), while male-sterility was observed in more variable field conditions (Vernoud et al. 2009). A second striking phenotype of *ocl4-1* mutants is the emergence of ectopic macrohairs in novel zones of the leaf, supporting a hypothesis that OCL4 is required to repress this epidermal structure in specific leaf domains.

The 0.4 mm size of maize anthers is notable for the high accumulation of 21-nt, phased, small interfering RNAs (phasiRNAs) that peak in abundance at this pre-meiotic stage (Zhai et al. 2015). First described in the grasses (Johnson et al. 2009), male reproductive tissues of diverse angiosperms accumulate two size classes of 21- and 24-nucleotide (nt) phasiRNAs (Zheng et al. 2015; Xia et al. 2019). The similarity of phasiRNAs and mammalian piRNAs (*P*-element induced wimpy testis -interacting RNAs) may be a case of convergent evolution in support of male reproduction (Zhai et al. 2015). The shared characteristics include a high abundance in male reproductive organs, two distinct size classes, lack of target mRNAs, phased production from non-coding precursors, and generation in somatic cells. Even as the precise direct functions of these RNA species remain unknown, the absence or perturbation of mammalian piRNAs or grass phasiRNAs is often associated with male-sterility.

Plant phasiRNAs originate from 5’-capped and polyadenylated, noncoding mRNA precursors (*PHAS* transcripts) that are generated by RNA polymerase II (Pol II). The precursors are derived from non-repetitive genomic loci. Precursor processing is initiated by the binding of a microRNA (miRNA) “trigger” : miR2118 for 21-nt *PHAS* loci and miR2275 for 24-nt *PHAS* loci. miRNA binding is required for Argonaute-mediated cleavage of the precursor, and the resulting 3’ fragment is converted to a double-stranded substrate by RNA-DEPENDENT RNA POLYMERASE 6 (RDR 6). The dsRNA is then cleaved at precise intervals by a Dicer-like enzyme: DCL4 generates 21-nt phasiRNAs, and DCL5 produces the 24-nt class. OCL4 is a master regulator of 21-nt phasiRNA biogenesis: it is required for robust production of *PHAS* precursor transcripts, normal levels of miR2118, and accumulation of 21-nt phasiRNAs (Zhai et al. 2015).

Nuclear-controlled, environment-conditioned, genic male sterility (EGMS) is a trait exploited by plant breeders, most successfully in rice. The molecular basis of this trait in rice has only recently been determined (Kim and Zhang 2018).The underlying mechanisms and players controlling this trait mainly include epigenetic regulation, transcription factors and non-coding RNAs. The rice PA64 male sterile line exhibits higher DNA methylation levels under long-day and high-temperature conditions (Chen et al. 2014), whereas small RNAs, long-non-coding RNAs and/or RNA-directed DNA methylation plays a role in regulating photoperiod-sensitive male sterility in Nongken 58S rice (Ding et al. 2012a, 2012b; Kim and Zhang 2018; Fan et al. 2016).The temperature-sensitive male sterility in the HengnongS-1 rice genotype results from a recessive single nucleotide polymorphism (SNP) within *PERSISTANT TAPETAL CELL 1*(*PTC1*), which encodes a PHD-finger protein involved in tapetal cell death and pollen development (Li et al. 2011; Qi et al. 2014). A mutation in the R2R3 MYB transcription factor, CARBON STARVED ANTHER, shows complete male sterility under short-day conditions, but is fertile under long-day conditions (Zhang et al. 2010; Zhang et al. 2013). Further, UPD-glucose pyrophosphorylase 1 (*Ugp-1*) co-suppression rice lines are male sterile under normal temperatures because endogenous *Ugp1* transcripts are not spliced, but revert to being male-fertile at low temperatures because of more efficient splicing of the primary *Ugp1* mRNA (Chen et al. 2007). One additional example is from the wheat line BS366, which is male-sterile at 10°C and fertile at 20°C, shows differential expression of miRNAs involved in modulating auxin and gibberellin signaling pathways for male fertility transition (Bai et al. 2017). It is clear that there are multiple pathways in which mutations can yield environment-conditioned male fertility.

Of particular relevance to our work are EGMS traits involving 21-nt phasiRNAs and/or long non-coding RNAs (lncRNAs). Photoperiod-sensitive male sterility in Nongken 58S rice is controlled in part by a 21-nt phasiRNA-producing locus (*Pms1*) that generates the *PMS1T* precursor transcript. *PMS1T* has a miR2118 target site and is processed to form abundant 21-nt phasiRNAs. Nongken 58S has a SNP in *PMS1T* near the miR2118 recognition site, and under long-day photoperiod conditions, this genotype is male sterile (Fan et al. 2016). A second locus, *Pms3* encodes a lncRNA that is also processed into small RNAs (Ding et al. 2012a; Zhou et al. 2012). A SNP in *pms3* promotes increased siRNA-directed methylation within its promoter region, leading to reduced transcript levels and consequent sterility, specifically under long-day conditions (Ding et al. 2012a; Zhou et al. 2012).

To date, no individual 21-nt *PHAS* locus has been associated with male sterility in maize. *ocl4* mutants lack nearly all 21-nt phasiRNAs (Zhai et al. 2015). *dcl5* mutants lack most 24-nt phasiRNAs and are male sterile under optimal, warm growing conditions (28°C/ 22°C, 16 hour days/ 8 hour nights in the greenhouse). Under cool growing conditions, fertility is restored in the absence of 24-nt phasiRNAs. Consequently, *dcl5* represents a case of EGMS. *Dcl5* is preferentially present in the tapetum; *dcl5* mutants show delayed tapetal development, starting in early meiosis (Teng et al. 2020). Combining our current knowledge about rice *pms1, pms3*, and maize *dcl5*, we suggest that reproductive phasiRNAs buffer development under variable or unfavorable environmental conditions.

Here, we describe an *ocl4//ocl4* derivative line, in which fertility was restored and perpetuated over several generations, under warm growing conditions. We designate this line as “conditionally fertile *ocl4*” as its fertility is apparently conditioned by ambient temperature. We present a genetic, phenotypic, RNA-seq, and small RNA-seq assessment of male-sterile *ocl4* and conditionally fertile *ocl4* anthers, in an effort to better understand the role of 21-nt phasiRNAs in male reproduction. Our results indicate that conditional male fertility of *ocl4* is associated with enhanced 21-nt phasiRNA accumulation in pre-meiotic maize anthers, while the anthers retained all canonical features of sterile *ocl4* cellular organization.

## Results

### *Ocl4* maintains plant fertility under cooler conditions

The *ocl4-1* allele in the A188 background (87.5% A188) was a gift of V. Vernoud prior to publication (Vernoud et al. 2009) for use in allelism testing of a panel of 55 new cases of recessive male sterility (Timofejeva et al. 2013). A line segregating 1:1 fertile:sterile was established and maintained by male-sterile (*ocl4//ocl4*) × heterozygous fertile (*ocl4//+*) crosses within a family for three field generations. Both the sterile and fertile individuals were used in complementation crosses to test for allelism in the new panel of male-sterile mutants. After the test cross, F_1_ progeny were observed for sterility in the subsequent summer field. In complemented families (no male sterility), several self-pollinations were performed to derive double mutant stocks. No unusual segregation ratios were observed in crosses involving *ocl4-1*over the course of several years (Timofejeva et al. 2013).

Because we observed excess male-sterility in five complementation tests involving a new, uncloned mutant, *tapetal cell layers 1*(*tcl1*), the F_2_ progenies of its complementation tests were re-evaluated in the greenhouse, including one family from the cross *ocl4//ocl4* × *tcl1/+*, followed by self-pollination. One set of progeny were grown in the greenhouse and genotyped by polymerase chain reaction (PCR) (Vernoud et al. 2009); two homozygous *ocl4* progeny were male-sterile as expected but two were fully fertile. The fertile individuals were successfully self-pollinated and crossed onto *ocl4* male-sterile ears; all progeny should have been *ocl4* homozygotes. As shown in Table 1, in the 2014 field season, seeds of one selfed individual were planted as ZM68, and 80% of the progeny were fertile (24F:6ms); each plant in this family was test-crossed to *tcl1//tcl1* male-sterile plants to identify individuals that were not carrying *tcl1*. Individual ZM68-10 was not a carrier and its self-pollination progeny and progeny from a cross onto sibling male-sterile ZM68-2 were selected for further analysis in subsequent field seasons. All plants were genotyped to confirm that they were *ocl4//ocl4*, and this was checked for at least two generations. Progeny of ZM68-10 (×) were 36F:0ms in the 2015 field season, and the ZM68-2ms × ZM68-10 progeny were 28F:17ms. These results indicate that the switch to fertility can persist, but it is not a complete switch to full fertility. We wondered if it reflected a genetic change in the ZM68-10 lineage.

**Table 1.**
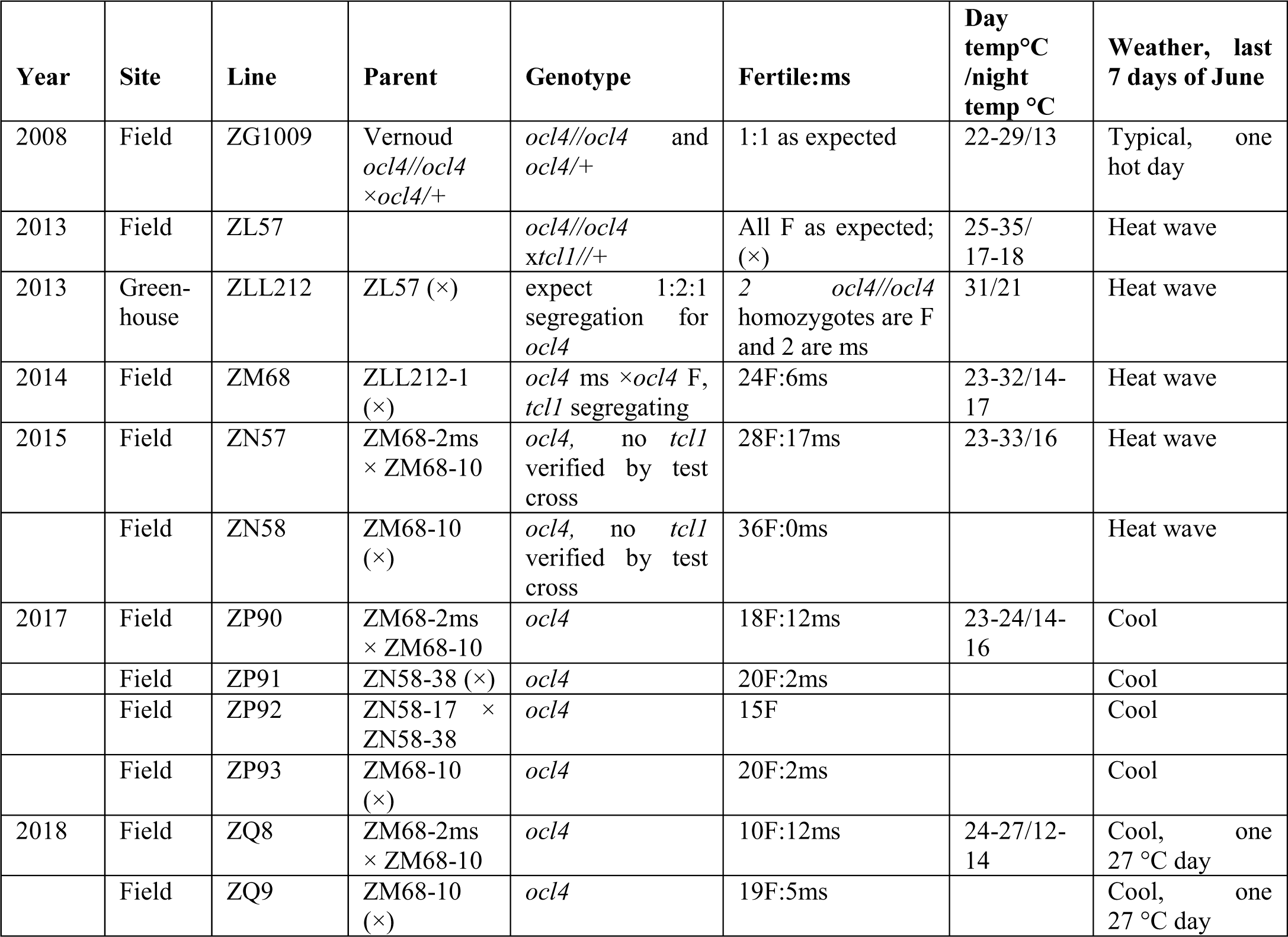
Derivation and propagation of fertile *ocl4* homozygous plants with fertility scoring.

| Year | Site | Line | Parent | Genotype | Fertile:ms | Day temp °C /night temp °C | Weather, last 7 days of June |
| --- | --- | --- | --- | --- | --- | --- | --- |
| 2008 | Field | ZG1009 | Vernoud<br>*ocl4//ocl4*<br>×*ocl4/+* | *ocl4//ocl4* and<br>*ocl4/+* | 1:1 as expected | 22-29/13 | Typical, one hot day |
| 2013 | Field | ZL57 |  | *ocl4//ocl4*<br>*xtcl1//+* | All F as expected;<br>(×) | 25-35/<br>17-18 | Heat wave |
| 2013 | Green-house | ZLL212 | ZL57 (×) | expect 1:2:1 segregation for<br>*ocl4* | 2 *ocl4//ocl4* homozygotes are F and 2 are ms | 31/21 | Heat wave |
| 2014 | Field | ZM68 | ZLL212-1 (×) | *ocl4* ms ×*ocl4* F, *tcl1* segregating | 24F:6ms | 23-32/14-17 | Heat wave |
| 2015 | Field | ZN57 | ZM68-2ms × ZM68-10 | *ocl4*, no *tcl1* verified by test cross | 28F:17ms | 23-33/16 | Heat wave |
|  | Field | ZN58 | ZM68-10 (×) | *ocl4*, no *tcl1* verified by test cross | 36F:0ms |  | Heat wave |
| 2017 | Field | ZP90 | ZM68-2ms × ZM68-10 | *ocl4* | 18F:12ms | 23-24/14-16 | Cool |
|  | Field | ZP91 | ZN58-38 (×) | *ocl4* | 20F:2ms |  | Cool |
|  | Field | ZP92 | ZN58-17 × ZN58-38 | *ocl4* | 15F |  | Cool |
|  | Field | ZP93 | ZM68-10 (×) | *ocl4* | 20F:2ms |  | Cool |
| 2018 | Field | ZQ8 | ZM68-2ms × ZM68-10 | *ocl4* | 10F:12ms | 24-27/12-14 | Cool, one 27 °C day |
|  | Field | ZQ9 | ZM68-10 (×) | *ocl4* | 19F:5ms |  | Cool, one 27 °C day |

We considered the possibility that we had selected a new *ocl4* allele. Evaluation of RNA-seq data of derivative progeny indicated that the original allele was still present (Online Resource 1 Fig. S1), as described in detail in the subsequent section.

We next considered environmental parameters. *ocl4-1* lines were planted in May each year, and the most advanced anthers in a tassel would reach the 1.25 mm stage (prophase I of meiosis) by the end of June, about 38 days after planting. Each tassel contains anthers spanning about 5 days of development in the A188 background; consequently the youngest anthers in the tassel would be at the 0.4 mm stage in the same tassel. Stanford typically has cool late June growing conditions of 22 – 24 °C daytime and 12 – 15 °C night temperatures, punctuated by one day of much cooler or much hotter weather. As indicated in Table 1, both the 2014 and 2015 growing seasons experienced heat waves at the end of June, while subsequent years had the typical cool growing conditions present during the complementation crosses earlier. Under heat wave conditions, the ZM68-10 (×) progeny were all fertile (36F:0ms), but under cooler conditions, the families were 64F:9ms. For ZM68-2ms × ZM68-10, under heat wave conditions the ratio was 28F:17ms, but under cool conditions, 28F:24ms. These results are indicative of EGMS – that is, higher temperatures favor fertility. Additionally, there appeared to be an epigenetic component, in that the putative epialleles in the ZM68-10 plant were highly responsive to heat, whereas the allele from the ZM68-2 plant was non-responsive or less responsive. This is an unexpected complication to EGMS and could involve a change in a locus other than *ocl4*.

### Conditionally fertile *ocl4* anthers retain a partially duplicated endothecial layer and exhibit a range of fertility levels

The observed restoration of fertility motivated a more detailed investigation of the extent of fertility in *ocl4-1* sterile compared to *ocl4-1* conditionally fertile plants. To achieve this, we used a scoring system to assess plant fertility by comparing the number of anthers in *ocl4* plants relative to normal *Ocl4* plants in families segregating 1:1 fertile:male-sterile. Full fertility was assigned a score of 5, and complete sterility a score of 0. Conditionally fertile *ocl4//ocl4* plants exhibited a range of fertility levels from 1 to 4 in the 2018 field season (Fig. 1a). We ascribe the phenotypic range to responses of cohorts of developing anthers to temperature conditions at a phenocritical stage (Table 1). We used the material with graded fertility scores for genotyping, anther dissection, and RNA preparation as indicated in Table 2.

**Table 2.** Derivation of lines used for phenotypic scoring, molecular analysis, and confocal microscopy. Conditionally fertile *ocl4* lines all trace back to individual ZLL212-1 (Table 1). PCR was used to validate genotype as described by Vernoud et al. (2009). ZOO lines were grown in the greenhouse; ZO and ZQ lines were field-grown. P refers to the prior parental generations. *ocl4* lines are homozygous for the mutant allele. Standard lines are maintained by crossing *ocl4//ocl4* male-sterile plants by *Ocl4//ocl4* heterozygotes (1:1 segregation fertile:sterile progeny) or by selfing heterozygotes (3:1 segregation).

| Conditionally fertile *ocl4* lines |  |  |  |  |  |  |  |  |
| --- | --- | --- | --- | --- | --- | --- | --- | --- |
|  | PCR check | P1 | PCR check | P2 | PCR check | P3 | PCR check | Uses |
| ZOO16 | yes | ZN57-17 ms × ZN57-38 | yes | ZM68-10 (×) | yes | ZLL212 ms × ZLL212 *ocl4* | yes | Phenotype, RNA extraction |
| ZOO19 | yes | ZN57-17 ms × 57-38 F | yes | ZM68-10 (×) | yes | ZLL212 ms × ZLL212 *ocl4* | yes | Phenotype, RNA extraction |
| ZOO21 | yes | ZN58-17 (×) | yes | ZM68-10 (×) | yes | ZLL212 ms × ZLL212 *ocl4* | yes | Phenotype, RNA extraction |
| ZQ8 | no | ZM68-2 ms × ZM68-10 | yes |  |  |  |  | Phenotype, Confocal microscopy |
| Standard *ocl4* sterile lines |  |  |  |  |  |  |  |  |
| ZO14 | yes | ZL11-3 ms × ZL fertile | no |  |  |  |  | Phenotype, RNA extraction |
| ZQ4 | no | ZP12 (×) | no | three generations ms × F |  | ZLL212 ms × ZLL212 fertile *Ocl4* | yes | Phenotype, Confocal microscopy |

**Fig. 1.**
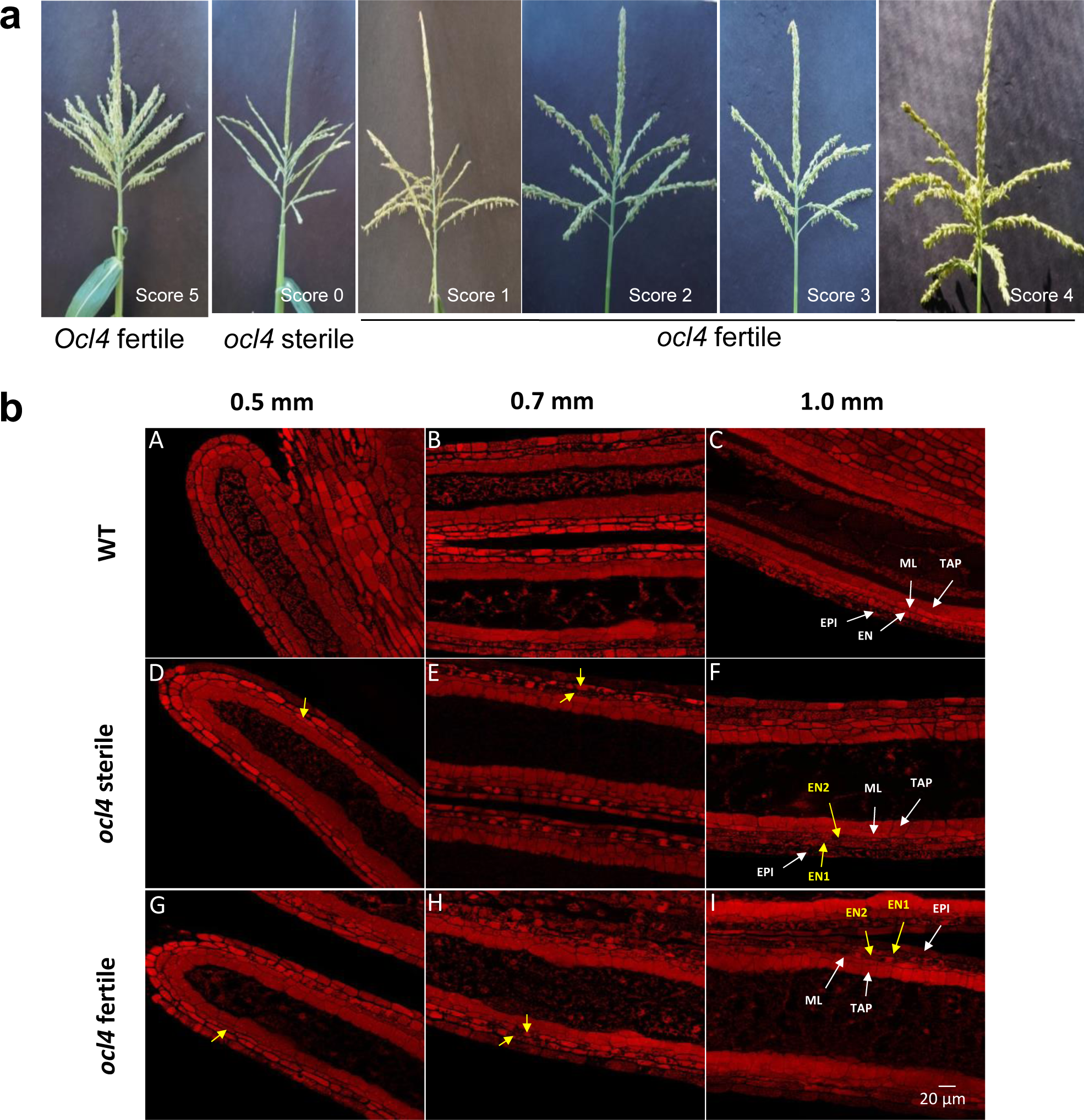
Tassel phenotype and anther cell wall organization in *ocl4* mutants. **(a)** Representative maize tassels at the pollen shedding stage from normal fertile (*Ocl4*) and *ocl4//ocl4* mutants, depicting sterile (*ocl4* sterile from a segregating 1:1 family) and the range of conditionally fertile (*ocl4* fertile) phenotypes. The scores represent fertility scores visually assigned on a scale of 0 to 5 (0= sterile; 5= wild type fertility). **(b)** Longitudinal confocal images of pre-meiotic anthers comparing fertile *Ocl4* and homozygous *ocl4//ocl4* mutants. Anthers of 0.5 mm, 0.7 mm, and 1.0 mm were dissected, fixed, stained with propidium iodide, and observed by confocal microscopy. Longitudinal confocal images of *Ocl4* fertile (labeled as WT) (A-C), *ocl4* sterile (D-F), and *ocl4* conditionally fertile (G-I) anthers at three developmental stages are presented. In *ocl4//ocl4* mutants, an extra cell layer with endothecial characteristics forms by periclinal division of the original endothecium (D-E, G-H; yellow arrowheads point to cells formed by extra periclinal division). Ultimately, *ocl4* lobes contain five somatic cell layers (F and I), in contrast to the four somatic cell layers found in normal *Ocl4* fertile lobes (A-C). Abbreviations: EPI, epidermis; EN, endothecium; ML, middle layer; TAP, tapetum; EN1 and EN2 represent two layers with endothecial characteristics. Scale bar: 20 μm.

The defining anatomical defect of the *ocl4* mutant is a partially duplicated endothecial layer in anther lobes. Using confocal microscopy, we found that the conditionally fertile *ocl4* retains this defect (Fig. 1b). In pre-meiotic *Ocl4* anthers, the expected architecture of a three-layer cell wall (epidermis, endothecium, secondary parietal layer) is present at 0.5 mm, but by 0.7 mm and completing by 1.0 mm, the normal four-layer cell wall exists after periclinal division of the secondary parietal cells to form the middle layer and tapetum. In contrast, in both *ocl4* sterile and *ocl4* conditionally fertile mutants, two cell layers with endothecial cell shape characteristics were gradually generated after extra periclinal cell divisions in the presumptive endothecial layer, resulting in five somatic cell layers by 1.0 mm (Fig. 1b) in the distal hemisphere of each lobe as previously described (Vernoud et al. 2009; Wang et al. 2012).

Previously, this extra subepidermal cell layer has been cited as the cause of pollen failure in *ocl4* mutants. Our findings question this link between a partial extra endothecial layer and male sterility. We cannot be definitive, however, because at the time of anther analysis, it is unknown whether a particular anther will be fertile or sterile. As shown in Fig.1a, there is a range of partial fertility in *ocl4* homozygous plants; it is possible that the confocal microscopy was of anthers destined to be sterile.

### Conditionally fertile *ocl4* anthers show higher accumulation of 21-nt phasiRNAs

Our observation of conditionally fertile *ocl4* anthers motivated us to assess the impact of *ocl4* on smallRNAs in general, and phasiRNAs in particular, given their known connection to male sterility. We built libraries and sequenced samples from *ocl4* conditionally fertile anthers at pre-meiotic stages (anthers length ranging from 0.4 to 0.7 mm) to compare with previously published *ocl4* sterile and wild type samples (Online Resource 2 Table S1). We first looked at the populations of different small RNA classes categorized by length to determine classes that are predominant and that are significantly impacted in the two *ocl4* conditions. We found the 21- and 24-nt small RNA classes to be the predominantclasses across all samples. Unlike the 24-nt class, we saw a significant reduction in small RNA abundance for the 21-nt class in both *ocl4* sterile and conditionally fertile samples compared to the wild type samples (Fig. 2a). *Ocl4* is a master regulator of 21-nt phasiRNAs; when absent, phasiRNAs are almost negligible (Zhai et al. 2015). To find out whether phasiRNAs are also negligible in the conditionally fertile *ocl4* anthers, we selected the 359 most abundant phasiRNA precursors (from 359 *PHAS* loci) with a predicted miR2118 complementary site, from the previously published list (Zhai et al. 2015) to examine the proportion of 21-nt phasiRNAs among the 21-nt small RNA class across samples. The accumulation levels and the fraction of phasiRNAs were the highest in wild type as expected, but were also significantly higher in the conditionally fertile *ocl4* compared to *ocl4* sterile samples, suggesting a partial restoration of phasiRNAs in conditionally fertile plants (Fig. 2b).

**Fig. 2.**
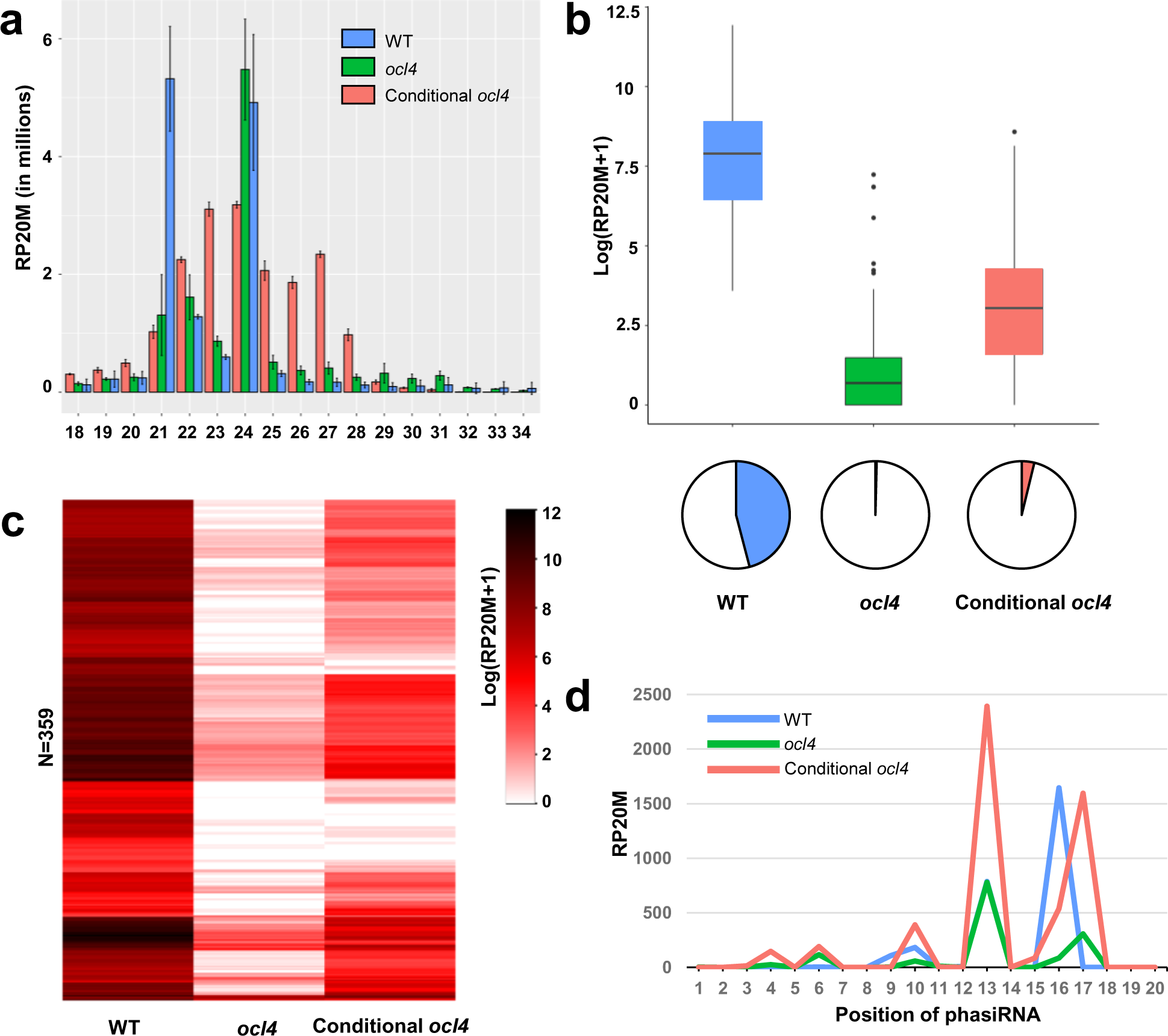
Conditionally fertile *ocl4* plants show relatively higher accumulation of phasiRNAs compared to *ocl4* sterile plants. **(a)** Bar plot showing abundance distribution of different size classes (18 to 34 nt in length) of small RNAs in wild type, *ocl4*, and conditionally fertile *ocl4* anthers. **(b)** Boxplot (top) displaying phasiRNA abundance distribution and pie chart (bottom) showing the fraction of phasiRNA abundance out of the total abundance of all the small RNA population. **(c)** Heatmap displaying the accumulation of phasiRNAs from 359 analyzed phasiRNA precursors. **(d)** Line plot illustrating a phasiRNA precursor which shows higher accumulation of phasiRNAs in conditionally fertile *ocl4* than the wild type anthers. RP20M stands for reads per 20 million reads. Samples used: for wild type (N=3) anthers of length 0.4 mm were used, for *ocl4* (N=2) and *ocl4* conditionally fertile (N=2) a mixture of anthers ranging from 0.4 and 0.7 mm in length were used.

We next explored phasiRNA accumulation from individual *PHAS* loci, as normally there is a wide variation in output. Out of 359 analyzed loci, most loci demonstrated negligible accumulation of phasiRNAs in *ocl4* mutants; however, there was a distinct subset of *PHAS* loci with significant accumulation of 21-nt phasiRNAs in conditionally fertile but not in *ocl4* sterile mutants (Fig. 2c). Although phasiRNAs are generated by sequential dicing of a *PHAS* locus transcript, there is unexpectedly non-stoichiometric yield of 21-nt products: typically one dominant (most abundant) phasiRNA is produced per locus (Tamim et al. 2018). Focusing on the dominant products, we found an example in which the conditionally fertile samples accumulated higher product than that the wild type samples. From the analysis at the phasiRNA level we observed the dominant phasiRNA (at position 13) to be highly abundant and three times higher in conditionally fertile samples (Fig. 2d). Therefore, while *Ocl4* is the master regulator of 21-nt phasiRNAs, warmer environmental conditions result into an increased accumulation of 21-nt phasiRNAs encoded by some loci.

### *Ago18b* is significantly up-regulated in conditionally fertile *ocl4* anthers

One possible explanation for the low yield of 21-nt phasiRNAs in *ocl4* mutants is low accumulation of the miR2118, the microRNA required for the initial cleavage of the primary transcripts. In a previous study of the *ms23* male-sterile maize mutant, there were few 24-nt phasiRNAs and miR2275 is nearly absent (Nan et al. 2017). Previously, low levels of miR2118 were found in sterile *ocl4* anthers (Zhai et al. 2015). These results implicate OCL4 as a major direct or indirect regulator of miR2118 biogenesis. In re-evaluating miR2118 in *ocl4*, we paradoxically found about a 4-fold reduction in *ocl4* sterile, but a 20-fold reduction in conditionally fertile *ocl4* anthers (Fig. 3a). Interestingly, in *Ocl4*, multiple miR2118 family isoforms accumulated, while in the conditionally fertile *ocl4* anthers, two-thirds of the overall abundance of the miR2118 family corresponded to just miR2118b (Fig. 3a). The miR2118b levels appear to be sufficient to generate 21-nt phasiRNAs. We next re-evaluated the impact of *ocl4* on the accumulation of transcripts from the 359 21-nt *PHAS* loci whose phasiRNA products were analyzed in the previous section. We confirmed that the *ocl4* mutation caused significant reduction in the overall abundance of *PHAS* precursor transcripts (Zhai et al. 2015). This reduction was less pronounced in conditionally fertile *ocl4* as compared to sterile *ocl4*. Further, at the level of individual loci, there were clear differences in accumulation between the sterile and fertile versions of the *ocl4* mutants (Fig. 3b; Online Resource 1 Fig. S2).

**Fig. 3.**
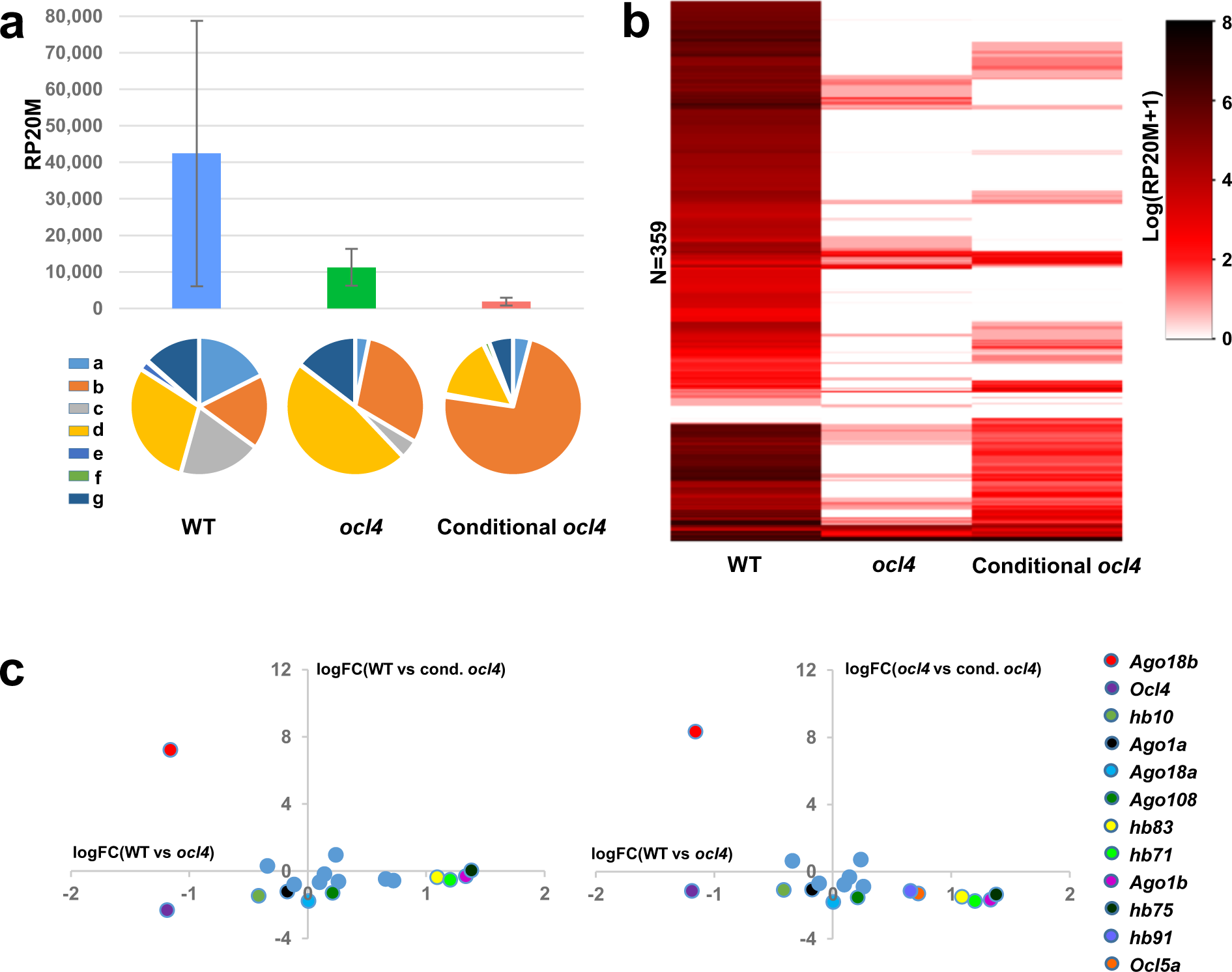
*Ocl4* controls key players that are upstream or downstream of the produced phasiRNAs. **(a)** Bar plot (top) showing total abundance of miR2118 family and pie chart (bottom) showing the fraction contribution of each family member between wild type, *ocl4*, and conditionally fertile *ocl4* anthers. Characters a, b, c, d, e, f, g represents miR2118 members: miR2118a, miR2118b etc. **(b)** Heatmap displaying the expression of 359 phasiRNA precursors. **(c)** Dot plot highlighting differential expression of key phasiRNA biogenesis pathway genes and some of the homeodomain leucine zipper type IV (HD-ZIP IV) tassel specific genes. Genes that are significantly differentially expressed using a log fold change cutoff of 1 are displayed with a unique color while the rest are displayed by a blue color.

To extend our investigation of changes in factors that may be involved in 21-nt phasiRNA activities, we directed our focus to two additional aspects of the phasiRNA pathway. First, as noted previously, nine family members of HD-ZIP IV transcription factors, *Ocl4* included, are expressed in immature tassels. We hypothesized that in *ocl4* samples, one or a combination of the remaining eight factors could act as a substitute. Second, if there are any “functional” phasiRNAs produced in the conditionally fertile *ocl4* samples, we would expect to see expression of Argonaute proteins (Vazquez et al. 2004) involved in the loading of phasiRNAs. To address these hypotheses, we performed a pair-wise differential expression analysis between wild type, *ocl4* sterile, and *ocl4* conditionally fertile samples. Specifically, we focused on *Ocl4* and its eight relatives, nine *Ago* members (*Ago1a, b, c, d*/ *Ago5a, b, c*/ *Ago18a, b*), and two *Dcl* loci (*Dcl4* and *Dcl5*). Overall, we found *Ago18b* to be the most significantly differentially expressed gene with about 8 log fold increase in the *ocl4* conditionally fertile compared to the sterile samples. Moreover, we found seven of the nine HD-ZIP IV transcription factors (*Ocl4* included) to be differentially expressed, however, together with *Ocl4*, all were down-regulated in the *ocl4* samples suggesting that none of these was acting as a substitute for *Ocl4* (Fig. 3c). Both *Dcl4* and *Dcl5* appear to be expressed relatively similarly across the samples and did not meet the cutoff as significant differentially expressed genes.

### Differential expression analysis uncovers distinct profile of genes for plastid differentiation and heat stress response

Is the overall transcriptome impacted differentially in conditionally fertile *ocl4* anthers compared to the *ocl4* sterile tissue? Evidence of a global impact on transcript accumulation in the conditionally fertile case would strengthen our view that these anthers are substantially different. We analyzed the RNA-seq reads from *Ocl4* and from the two classes of *ocl4* 0.4 – 0.7 mm anthers: most genes were expressed in all three conditions (Fig. 4a). While there is a huge overlap (∼5000) between genes that are differentially expressed between *ocl4* sterile and *ocl4* conditionally fertile samples relative to the wild type, there are about ∼1500 genes that are uniquely differentially expressed in either *ocl4* or conditionally fertile *ocl4* samples. Furthermore, the presence of chloroplasts is a key characteristic of normal endothecial cells by the 1.0 mm anther stage accompanied by expression of nuclear genes involved in plastid differentiation (Murphy et al. 2015). We analyzed expression of a set of 238 nuclear genes that produce proteins localized to plastids genes in *Ocl4* normal, sterile, and conditionally fertile pre-meiotic maize anthers; 57 genes were found to be significantly differentially expressed (Fig. 4b). To find out more about the overall differentially expressed genes in conditional fertility, we performed a Gene Ontology (GO) analysis to search for processes that these genes play a role in. Interestingly, we found genes to be significantly enriched in processes that respond to heat as well as processes linked to the RNA Pol II regulation of transcription in response to stress (Fig. 4c; Online Resource 2 Tables S2 a, b, c).

**Fig. 4.**
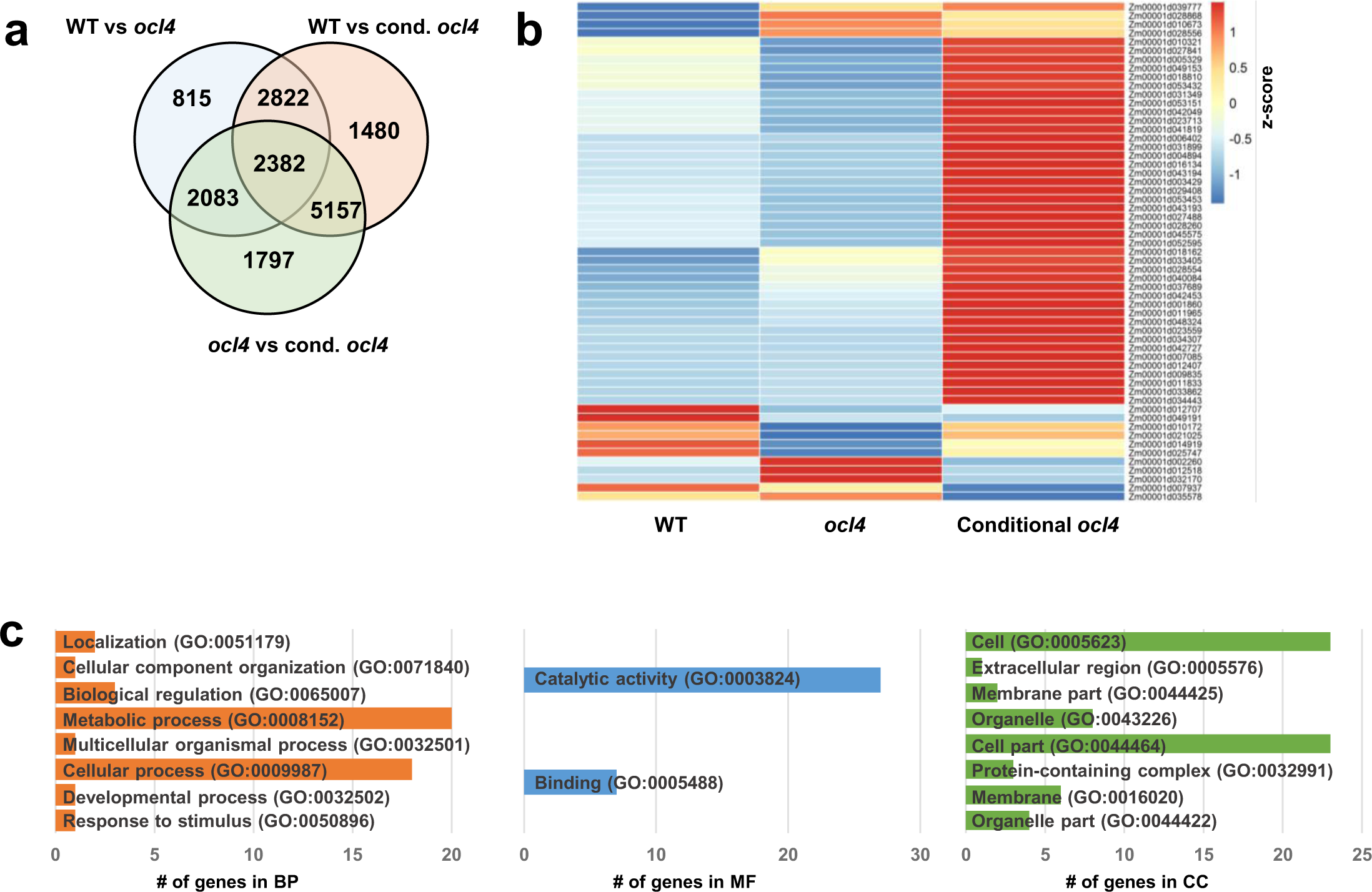
Conditionally fertile *ocl4* anthers have distinct profile of genes for plastid differentiation and heat stress response. **(a)** Venn diagram showing the overlap of genes that are differentially expressed between three pair-wise differential expression analysis between wild type, *ocl4*, and conditionally fertile *ocl4* samples. **(b)** Heatmap highlighting 57 differentially expressed genes from the set of 238 previously published nuclear genes involved in plastid differentiation. **(c)** Gene ontology (GO) terms and the respective list of genes that are enriched in Biological Process (BP), Molecular Function (MF), and Cellular Component (CC).

## Discussion

We used fertility classification, microscopy, genetics, and transcriptional analysis to characterize the maize *ocl4* mutant in which conditional male fertility can occur. Conditional fertility can be ascribed, at least in part, to a positive response to elevated temperature (Table 1). Collectively, the data presented here and prior observations (Zhai et al. 2015) demonstrate that the 21-nt class of phasiRNAs peaks in abundance in 0.4 mm anthers, during the loss of pluripotency in the newly specified somatic cells. These small RNAs are highly abundant through the 1.0 mm stage, when their precursor transcripts and the trigger miR2118 have decreased substantially. This suggests that there is a narrow developmental window during which 21-nt phasiRNAs are generated, and that they have a relatively long lifespan. We observed that the biogenesis of 21-nt phasiRNAs is largely dependent on *Ocl4* at three key steps: production of *PHAS* precursor transcripts (Fig. 3b), expression of individual miR2118 loci (Fig. 3a), and accumulation of 21-nt phasiRNAs (Fig. 2c). All three sets of RNAs were greatly reduced in sterile pre-meiotic anthers of the *ocl4* mutant, confirming a previous report (Zhai et al. 2015). The total 21-nt small RNAs dropped by ∼46 and ∼38 percent in sterile and fertile *ocl4* anthers respectively (Fig. 2a). The major classes of the reduced-abundance small RNAs include miR2118 family members (Fig. 3a) and 21-nt phasiRNAs (Fig. 2b). Besides the small RNA classes specifically linked to the biogenesis of 21-nt phasiRNAs, miR159, miR408, miR168, and miR2275 accumulation levels were much higher in conditionally fertile *ocl4* (Online Resource 1 Fig. S3). These changes indicated that *OCL4* plays a major role in 21-nt phasiRNA biogenesis during the pre-meiotic stage of maize anther development.

Surprisingly, in sterile *ocl4* anthers, we observed a subset of *PHAS* loci that are expressed at approximately normal levels, plus there is a low level of miR2118 accumulation, and 21-nt phasiRNAs are still present. It therefore seems possible that a small proportion of 21-nt phasiRNA production is controlled by a different transcription factor than OCL4. This possibility is reinforced by analysis of the conditionally fertile *ocl4* anthers: there was a partial restoration of 21-nt phasiRNAs (Fig. 2c), heightened expression of miR2118b (Fig. 3c), and normal accumulation of some *PHAS* precursor types and ectopic expression of a unique locus (at position 13) (Fig. 2d).

Another explanation for lower 21-nt phasiRNA levels in *ocl4* is that the *21-PHAS* precursors are less efficiently processed into 21-nt phasiRNAs, because there is reduced abundance of miR2118 in fertile *ocl4* (Fig. 3a). The level of *Dcl4* transcripts was essentially the same in all genotypes (Fig. 3c), while the *21-PHAS* precursor transcripts were moderately increased in fertile *ocl4* (Fig. 3b). Furthermore, there was no significant change in the phasing pattern among the genotypes and phenotypes that we analyzed (Fig. 2d), indicating that no miRNA trigger other than miR2118 initiates 21-nt phasiRNA production. Our studies raise interesting questions for future analysis. Why is there higherexpression of *Ago1b* and less miR2118 in the fertile *ocl4*? Further chromatin immunoprecipitation (ChIP) studies will be required to answer such questions and to determine, for example, if increased AGO1b protein in sterile *ocl4* stabilized miR2118 and therefore “increased” the abundance.

Our data also confirmed the observation that 24-nt phasiRNA abundances in *ocl4* are largely unchanged relative to the sibling *Ocl4* fully fertile plants (Zhai et al. 2015). We found only moderate fluctuations in *ocl4*, 17% higher in sterile and 32% lower in fertile pre-meiotic anthers, and these levels correlated with *Dcl5* transcript levels (Fig. 3c). We conclude that *OCL4* is unlikely to directly function in the 24-nt phasiRNA pathway.

These results suggest that OCL4 is responsible for the regulation of most, but not all 21-nt phasiRNA biogenesis. We have evidence of normal or higher accumulation of specific 21-phasiRNAs, a miR2118 isoform, and 21-nt *PHAS* transcript in *ocl4* mutants. In the conditionally fertile *ocl4* plants, these features are elevated relative to the *ocl4* sterile individuals. Based on these observations, it is likely that the OCL4 transcription factor is responsible for regulating most loci involved in 21-nt phasiRNA biogenesis, but there is a subclass regulated independently. Furthermore, this subclass is up-regulated in conditionally male-fertile *ocl4* anthers. We tested the logical possibility that another HD-ZIP IV transcription factor could explain these phenomena but found there are no HD-ZIP IV transcription factors that are up-regulated in *ocl4* conditionally fertile samples. Interestingly, we found that *Ago18b* is significantly up-regulated in the conditionally fertile samples (Fig. 3c). It is known that AGO18b is highly expressed in maize tassels during meiosis and it negatively controls determinacy of spikelet meristems. AGO18b primarily binds to 21-nt phasiRNAs in the pre-meiotic tassels (Sun et al. 2019). Knockdown of *Ago18* results in male sterility and reduction in phasiRNAs in rice (Das et al. 2020). Thus, it is possible that restoration of 21-nt phasiRNAs in conditionally fertile *ocl4* might be due to AGO18b accumulation, which may be independently stabilizing a sub-class of 21-nt phasiRNAs.

A possibility of an epigenetic component to conditional male fertility mediated by *ocl4* remains an open question for future analysis. It is also possible that parental environment can establish epigenetic states that influence multiple generations (Heard and Martienssen 2014). Environmentally-induced *trans*-generational epigenetic inheritance is an emerging field (Casier et al. 2019, Pang et al. 2019). The scientific understanding of myriad mechanisms, especially the role of reproductive small RNAs in *trans*-generational inheritance of male fertility and sterility, is at a nascent stage. In the nematode, *C. elegans* piRNAs are capable of triggering a multigenerational epigenetic memory in the germline (Ashe et al. 2012). Yet, studies implicating reproductive small RNAs in environmentally regulated or ‘conditional’ fertility are absent in mammals and rare in plants. Future studies focused on epigenetic factors and histone and DNA epigenetic marks using *ocl4* plants grown under controlled conditions could test the hypothesis that parental growth environment establishes the propensity for full sterility or partial fertility of *ocl4* mutants.

## Materials and methods

### Plant materials and growing conditions

V. Vernoud donated an *ocl4* stock introgressed in inbred A188 (Vernoud et al. 2009), and I. Golubovskaya (UC Berkeley) derived a stock segregating 1:1 for *ocl4* (*ocl4//ocl4* male sterile × *ocl4//+* maintainer) that was donated to Stanford University. This stock was maintained over three generations with 1:1 segregation, and used in 55 complementation tests with newly identified male-sterile mutants (Timofejeva et al. 2013). In 2013, the test cross with the *tcl1* mutant, in which the *ocl4//ocl4* male sterile female was crossed by a heterozygous *tcl1//+* male, was field-grown as family ZL57; all progeny were fertile, as expected, and self-pollination was performed to generate families expected to segregate 1:2:1 for *ocl4*. The first instance of fertility in an *ocl4* homozygote was in the greenhouse generation of 2013 (ZLL212 family). At the “ZLL” generation, greenhouse-grown plants were genotyped using a published PCR-based protocol (Vernoud et al. 2009), and two homozygous *ocl4//ocl4* fertile plants were identified (Table 1). Progeny of these individuals were grown in subsequent field seasons, genotyped again at most generations to verify homozygosity for *ocl4*. Some field seasons experienced late June heat waves corresponding to the time period of early anther development with abundant 21-nt phasiRNAs. Heat wave conditions correspond to three or more days in a row of 32 °C or higher; night temperatures on such days are also higher than normal by several degrees. Historic weather data were retrieved from the Weather Underground dataset (https://www.wunderground.com/history) using the station closest (2 km) from the Stanford field site (37°25’45.9” N, 122°10’52.5” W).

### Anther collections and confocal microscopy

Anthers were dissected and staged by length using a micrometer and dissecting microscope. For RNA extraction, a total of 50 anthers of the target length (0.4 to 0.7 mm) were pooled from one immature tassel of each of the conditionally fertile *ocl4* plant. All the anthers were transferred to chilled vials immersed in liquid nitrogen and stored at -80°C until further use. For confocal microscopy, anthers of 0.5 mm, 0.7 mm, and 1.0 mm were dissected and stored in 100% ethanol at 4 °C. Propidium iodide staining and confocal imaging utilized published protocols (Kelliher and Walbot 2011).

### sRNA-seq and RNA-seq library construction and sequencing

Total RNA for sRNA-seq and RNA-seq libraries was isolated using the TRI Reagent RNA isolation reagent (Sigma-Aldrich) following the manufacturer’s instructions. Total RNA quality was assessed using the Agilent RNA 6000 Pico Kit (Agilent). For library preparation, 20 to 30 nt RNAs were excised from a 15% polyacrylamide/urea gel, and ∼25 ng of sRNA was used for library construction with the TruSeq Small RNA Prep Kit (Illumina) following the manufacturer’s instructions. For RNA-seq, 2 µg of total RNA was treated with DNase I (New England BioLabs) and then cleaned with RNA Clean and Concentrator-5 (Zymo Research). The TruSeq Stranded Total RNA with RiboZero-Plant Kit (Illumina) or NEBNext Ultra II Directional RNA Library Prep Kit (New England BioLabs) was used for library construction with 500 ng of treated RNA, following the manufacturer’s instructions. Sequencing in single-end mode on an Illumina HiSeq 2500 (University of Delaware), yielded 51 nt reads for both sRNA-seq and RNA-seq.

### Data handling and bioinformatics

sRNA-seq data were processed as previously described (Mathioni et al. 2017). Briefly, we first used Trimmomatic (v0.32) (Bolger et al. 2014) to remove the linker adaptor sequences. Trimmed reads are then chopped to reads that are within 18 to 34 nt length before they were mapped to version 4 of the B73 maize genome using Bowtie (Langmead et al. 2009). Read counts were normalized to 20 million to allow for the direct comparison across libraries and were hit normalized to account for reads that map in multiple positions in the genome. For phasiRNA analysis, we used the list of 463 previously reported *PHAS* loci (Zhai et al. 2015) and selected 359 *PHAS* loci with a miR2118 target site. For RNA-seq libraries, reads were first trimmed for adapter using Trimmomatic and then mapped to version 4 of the B73 genome using Tophat v2 (Trapnell et al. 2009). Read counts were then generated using featureCounts (Liao et al. 2014), and the differential expression analyses were performed using DESeq2 (Love et al. 2014) package in R.

## Supporting information

Supplemental Resource 1

Supplemental Resource 2

## Author contributions

VW conceived of the project and derived line(s) reported here, which were genotyped by HZ. PY collected staged anthers for RNA analysis and evaluated fertility status; XZ acquired confocal images; CT made libraries and ST analyzed sequencing reads. PY and ST interpreted the data in consultation with VW and wrote the manuscript, which was improved and edited by VW and BCM. All authors provided critical feedback and helped shape the research, analysis, and final manuscript.

## Acknowledgments

This study was supported by U.S. National Science Foundation Plant Genome Research Project (Award # 1754097) to BCM and VW. PY was supported by a Fulbright-Nehru grant from United-States India Educational Foundation (Award No. 2200/FNPDR/2016).

## Data deposition

The data reported in this paper have been deposited in the Gene Expression Omnibus (GEO) database, Series GSE150446, accession nos. GSM4550675 to GSM4550682 for RNA-seq data, GSM4550683 to GSM4550690 for small RNA data.

