## Supplemental Resource 1 for "Environment-conditioned male fertility of HD-ZIP IV transcription factor mutant *ocl4*: impact on 21-nt phasiRNA accumulation in pre-meiotic maize anthers"

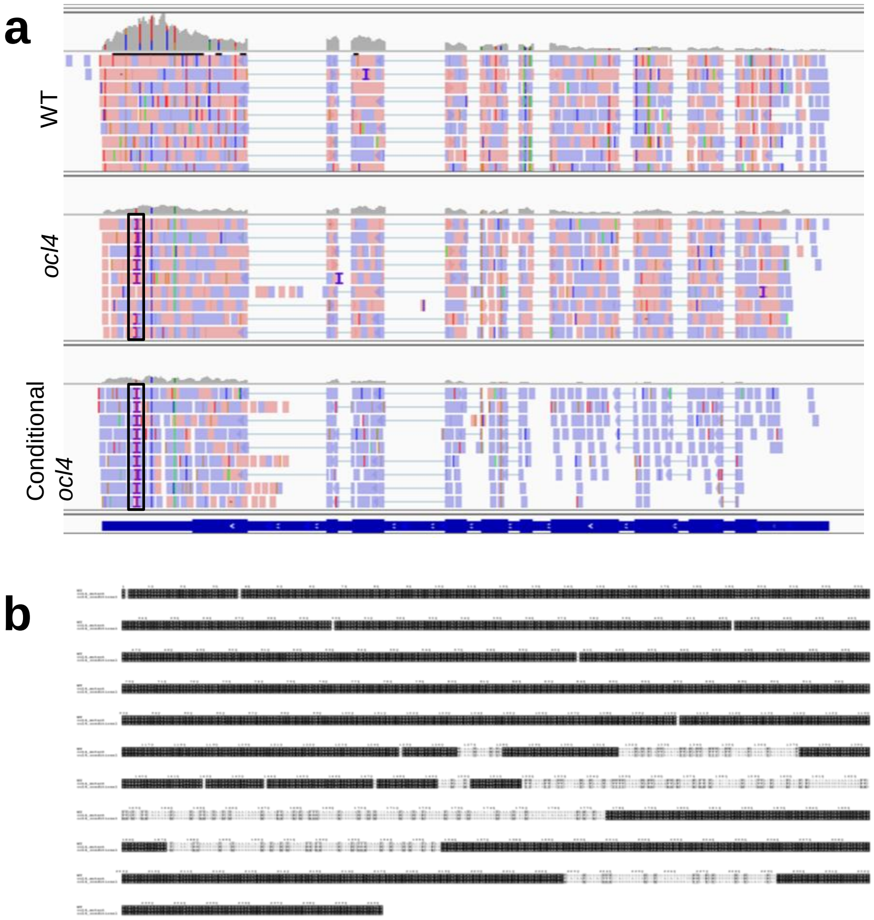


**Fig. S1 *ocl4* allele is maintained in the conditionally fertile *ocl4* samples. (a)** Alignment of RNA-seq reads from wild type, *ocl4*, and conditional *ocl4* along *Ocl4*. Regions along *ocl4* and conditional *ocl4* tracks highlighted by a black box correspond to the regions of *Mu8* insertion in exon 6. **(b)** Multiple sequence alignment of consensus reads obtained from the aligned reads in part a.

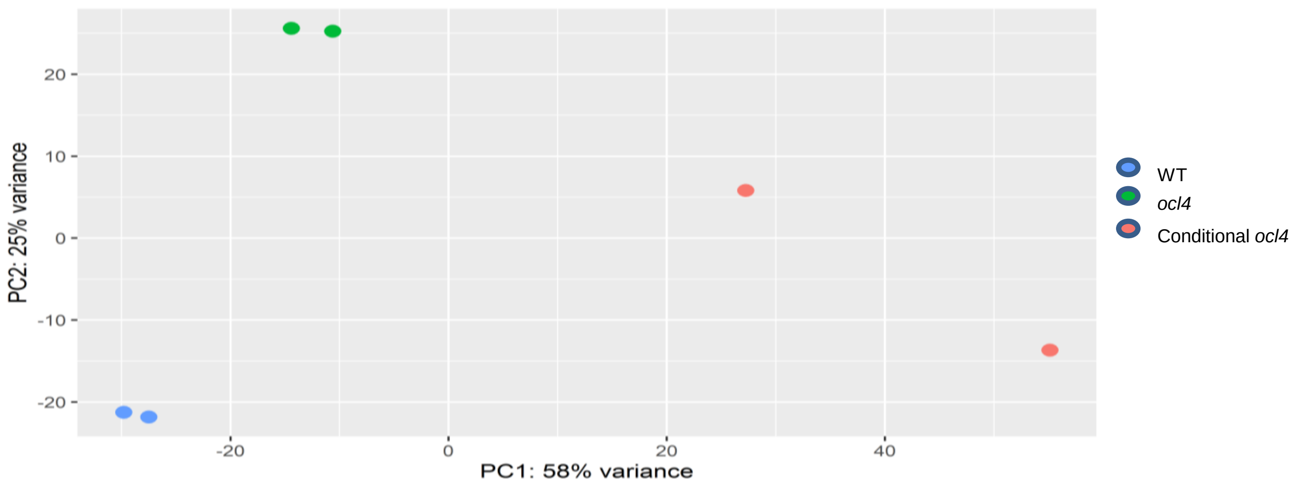


**Fig. S2 Wild type, *ocl4*, and conditional *ocl4* samples show strong variation.** PCA plot showing clustering of wild type, *ocl4*, and conditional *ocl4* samples

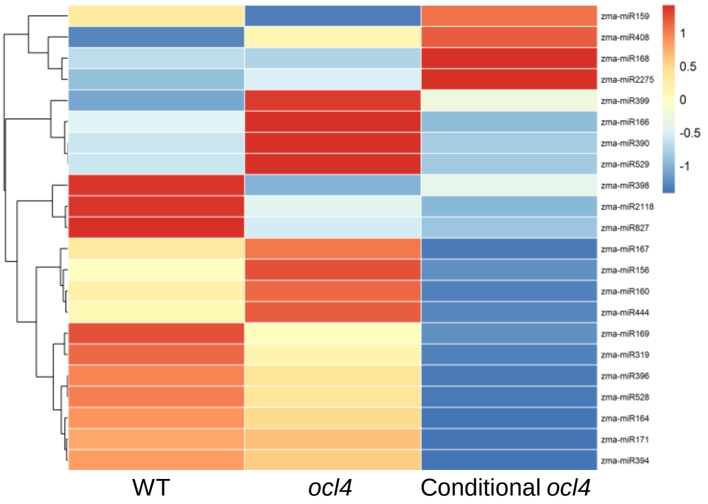


**Fig. S3 MicroRNA families impacted by *ocl4.*** Heatmap showing microRNA families that show significant accumulation difference between wild type, *ocl4,* and conditionally fertile *ocl4* samples*.* Abundances of each microRNA family was calculated as sum abundance of the respective family members.
