## Supplemental Resource 2 for "Environment-conditioned male fertility of HD-ZIP IV transcription factor mutant *ocl4*: impact on 21-nt phasiRNA accumulation in pre-meiotic maize anthers"

### Table S1 Summary of sRNS-seq and RNA-seq libraries

#### A. sRNA libraries

| Stage | Genotype | Male Fertility | Total Sequences [a] | Genome Matched Reads [b] | Distinct Genome Matched Reads[c] | t/r/sn/snoRNA Matched Reads[b] | GEO accession[d] |
|---|---|---|---|---|---|---|---|
| 0.4 mm Anther | Fertile (W23) | Fully fertile | 35,286,904 | 25,027,015 | 4,008,890 | 2,170,440 | GSM1262525 |
| 0.4 mm Anther | Fertile (W23) | Fully fertile | 36,257,470 | 26,056,428 | 1,219,871 | 1,185,886 | GSM1262536 |
| 0.4 mm Anther | Fertile (W23) | Fully fertile | 25,908,576 | 17,954,720 | 4,304,661 | 722,325 | GSM1262536 |
| 0.4 mm Anther | *ocl4* | Male sterile | 55,691,540 | 39,957,660 | 5,101,990 | 6,659,714 | GSM1262576 |
| 0.4 mm Anther | *ocl4* | Male sterile | 21,950,618 | 15,458,497 | 4,308,442 | 1,723,437 | GSM1262581 |
| 0.4-0.7 mm Anther | *ocl4* | Conditionally fertile | 19,664,132 | 9,360,710 | 1,160,799 | 6,765,795 | GSM4550684 |
| 0.4-0.7 mm Anther | *ocl4* | Conditionally fertile | 20,591,670 | 10,322,944 | 1,153,590 | 6,746,074 | GSM4550687 |

#### B. RNA-seq libraries

| Stage | Genotype | Male Fertility | Total Sequences [a] | Genome Matched Reads [b] | Distinct Genome Matched Reads[c] | t/r/sn/snoRNA Matched Reads[b] | GEO accession[d] |
|---|---|---|---|---|---|---|---|
| 0.4 mm Anther | Fertile (W23) | Fully fertile | 24,344,587 | 16,720,188 | 8,338,792 | 253,069 | GSM1262518 |
| 0.4 mm Anther | Fertile (W23) | Fully fertile | 21,258,204 | 12,157,106 | 6,860,089 | 28,375 | GSM4552958 |
| 0.4 mm Anther | ocl4 | Male sterile | 25,290,947 | 18,591,642 | 9,810,950 | 208,678 | GSM1262520 |
| 0.7 mm Anther | ocl4 | Male sterile | 27,271,637 | 19,857,673 | 10,083,208 | 507,477 | GSM1262521 |
| 0.4-0.7 mm Anther | ocl4 | Conditionally fertile | 32,139,342 | 20,104,170 | 7,250,879 | 584,949 | GSM4550676 |
| 0.4-0.7 mm Anther | ocl4 | Conditionally fertile | 32,380,429 | 20,340,334 | 6,810,001 | 422,838 | GSM4550679 |

[a]Total small RNAs after filtering bad reads and trimming adapters.
[b]Numbers determined by mapping to the maize B73 genome, version 4.
[c]Does not include the data listed in the column "t/rRNA Matched Reads".
[d]Libraries GSM1262525-1262582 are reused, previously published data (Zhai et al. 2015)

### Tables S2

#### a. List of genes represented in Biological Process (BP) represented by Gene Ontology (GO) category

| GO:0051179 | GO:0071840 | GO:0065007 | GO:0008152 | GO:0032501 | GO:0009987 | GO:0032502 | GO:0050896 |
|---|---|---|---|---|---|---|---|
| Zm00001d029408 | Zm00001d035578 | Zm00001d037689 | Zm00001d033405 | Zm00001d053453 | Zm00001d033405 | Zm00001d053453 | Zm00001d043194 |
| Zm00001d043194 | | Zm00001d031349 | Zm00001d049191 | | Zm00001d049191 | | |
| | | Zm00001d043194 | Zm00001d023559 | | Zm00001d023559 | | |
| | | | Zm00001d042727 | | Zm00001d042727 | | |
| | | | Zm00001d031899 | | Zm00001d031899 | | |
| | | | Zm00001d027488 | | Zm00001d012707 | | |
| | | | Zm00001d012707 | | Zm00001d048324 | | |
| | | | Zm00001d048324 | | Zm00001d027841 | | |
| | | | Zm00001d027841 | | Zm00001d040084 | | |
| | | | Zm00001d003429 | | Zm00001d035578 | | |
| | | | Zm00001d040084 | | Zm00001d010172 | | |
| | | | Zm00001d035578 | | Zm00001d037689 | | |
| | | | Zm00001d010172 | | Zm00001d031349 | | |
| | | | Zm00001d037689 | | Zm00001d012407 | | |
| | | | Zm00001d031349 | | Zm00001d012518 | | |
| | | | Zm00001d012407 | | Zm00001d034443 | | |
| | | | Zm00001d012518 | | Zm00001d043194 | | |
| | | | Zm00001d034443 | | Zm00001d009835 | | |
| | | | Zm00001d043194 | | | | |
| | | | Zm00001d009835 | | | | |

**b. List of genes represented in Molecular Function (MF) represented by Gene Ontology (GO) category**

| GO:0003824 | GO:0005488 |
|---|---|
| Zm00001d033405 | Zm00001d027488 |
| Zm00001d049191 | Zm00001d048324 |
| Zm00001d023559 | Zm00001d027841 |
| Zm00001d042727 | Zm00001d003429 |
| Zm00001d031899 | Zm00001d035578 |
| Zm00001d053453 | Zm00001d012518 |
| Zm00001d025747 | Zm00001d043194 |
| Zm00001d027488 | |
| Zm00001d012707 | |
| Zm00001d048324 | |
| Zm00001d053432 | |
| Zm00001d027841 | |
| Zm00001d016134 | |
| Zm00001d003429 | |
| Zm00001d040084 | |
| Zm00001d035578 | |
| Zm00001d010172 | |
| Zm00001d037689 | |
| Zm00001d002260 | |
| Zm00001d031349 | |
| Zm00001d012407 | |
| Zm00001d012518 | |
| Zm00001d034443 | |
| Zm00001d014919 | |
| Zm00001d043194 | |
| Zm00001d009835 | |
| Zm00001d007937 | |

### c. List of genes represented in Cellular Component (CC) represented by Gene Ontology (GO) category

| GO:0005623 | GO:0005576 | GO:0044425 | GO:0043226 | GO:0044464 | GO:0032991 | GO:0016020 | GO:0044422 |
|---|---|---|---|---|---|---|---|
| Zm00001d033405 | Zm00001d053453 | Zm00001d041819 | Zm00001d029408 | Zm00001d033405 | Zm00001d041819 | Zm00001d029408 | Zm00001d041819 |
| Zm00001d029408 | | Zm00001d023713 | Zm00001d053453 | Zm00001d029408 | Zm00001d035578 | Zm00001d041819 | Zm00001d023713 |
| Zm00001d049191 | | | Zm00001d041819 | Zm00001d049191 | Zm00001d023713 | Zm00001d053432 | Zm00001d018162 |
| Zm00001d023559 | | | Zm00001d023713 | Zm00001d023559 | | Zm00001d016134 | Zm00001d034443 |
| Zm00001d042727 | | | Zm00001d018162 | Zm00001d042727 | | Zm00001d023713 | |
| Zm00001d053453 | | | Zm00001d002260 | Zm00001d053453 | | Zm00001d018162 | |
| Zm00001d041819 | | | Zm00001d031349 | Zm00001d041819 | | | |
| Zm00001d048324 | | | Zm00001d034443 | Zm00001d048324 | | | |
| Zm00001d053432 | | | | Zm00001d053432 | | | |
| Zm00001d027841 | | | | Zm00001d027841 | | | |
| Zm00001d016134 | | | | Zm00001d016134 | | | |
| Zm00001d040084 | | | | Zm00001d040084 | | | |
| Zm00001d023713 | | | | Zm00001d023713 | | | |
| Zm00001d018162 | | | | Zm00001d018162 | | | |
| Zm00001d010172 | | | | Zm00001d010172 | | | |
| Zm00001d037689 | | | | Zm00001d037689 | | | |
| Zm00001d002260 | | | | Zm00001d002260 | | | |
| Zm00001d031349 | | | | Zm00001d031349 | | | |
| Zm00001d012407 | | | | Zm00001d012407 | | | |
| Zm00001d034443 | | | | Zm00001d034443 | | | |
| Zm00001d014919 | | | | Zm00001d014919 | | | |
| Zm00001d043194 | | | | Zm00001d043194 | | | |
| Zm00001d009835 | | | | Zm00001d009835 | | | |
